# Examining DNA Breathing with pyDNA-EPBD

**DOI:** 10.1101/2023.09.09.557010

**Authors:** Anowarul Kabir, Manish Bhattarai, Kim Ø. Rasmussen, Amarda Shehu, Anny Usheva, Alan R Bishop, Boian S Alexandrov

## Abstract

**Motivation:** The two strands of the DNA double helix locally and spontaneously separate and recombine in living cells due to the inherent thermal DNA motion.This dynamics results in transient openings in the double helix and is referred to as “DNA breathing” or “DNA bubbles.” The propensity to form local transient openings is important in a wide range of biological processes, such as transcription, replication, and transcription factors binding. However, the modeling and computer simulation of these phenomena, have remained a challenge due to the complex interplay of numerous factors, such as, temperature, salt content, DNA sequence, hydrogen bonding, base stacking, and others.

**Results:** We present pyDNA-EPBD, a parallel software implementation of the Extended Peyrard-Bishop-Dauxois (EPBD) nonlinear DNA model that allows us to describe some features of DNA dynamics in detail. The pyDNA-EPBD generates genomic scale profiles of average base-pair openings, base flipping probability,DNA bubble probability, and calculations of the characteristically dynamic length indicating the number of base pairs statistically significantly affected by a single point mutation using the Markov Chain Monte Carlo (MCMC) algorithm.

## 1 Introduction

The structural integrity of biological macromolecules is primarily governed by hydrogen bonds (H-bonds) Figure 1a), which have natural vibration frequencies in the terahertz range Turton et al. (2014). H-bonds are much weaker (∼ few *k* _*B*_ Ts) than covalent bonds, causing the macromolecules to experience slow conformational motion resulting from the inherent thermal fluctuations at physiological temperatures.

**Fig. 1:**
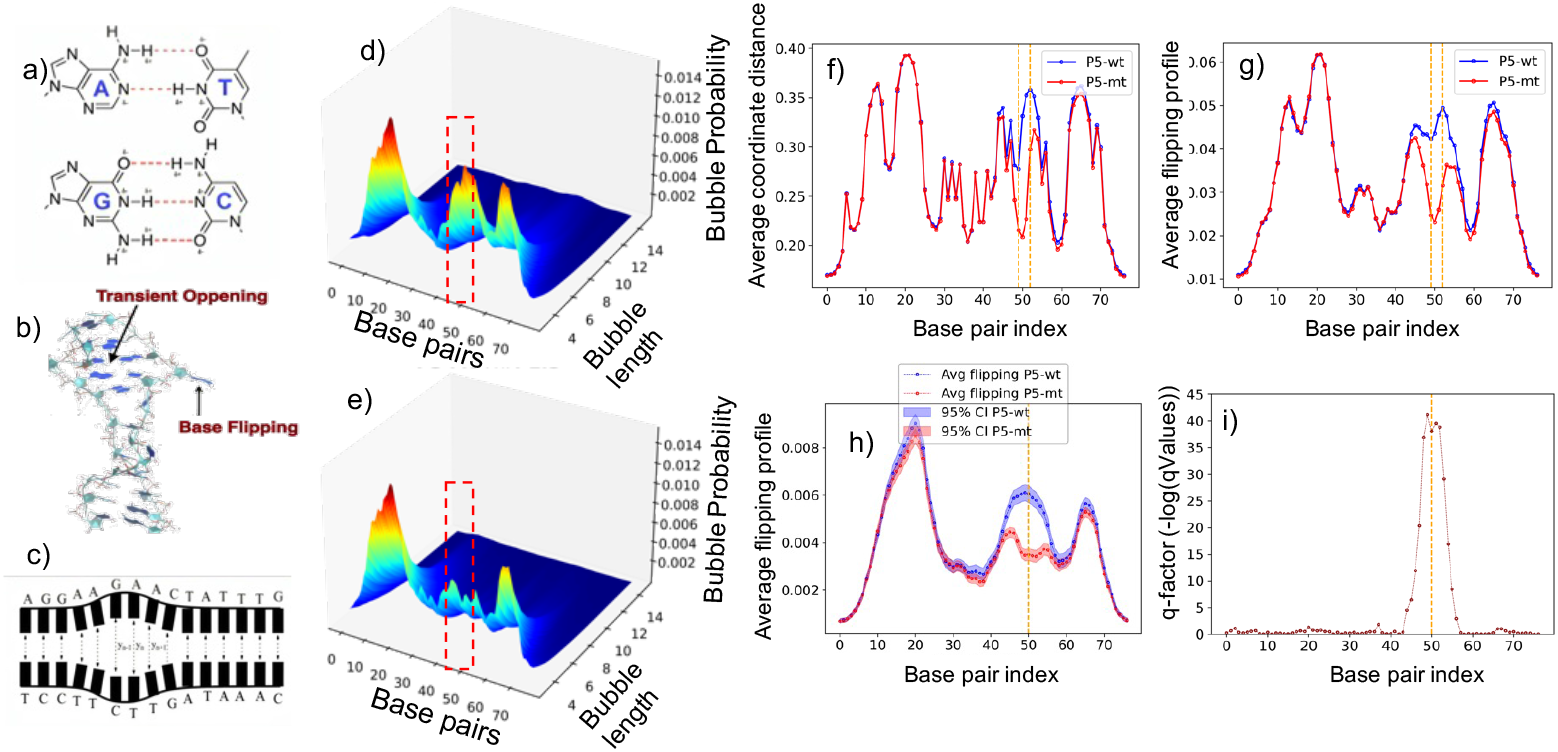
DNA Breathing Dynamics and Analysis. (a) The primary governance of macromolecules through hydrogen bonds (H-bonds). (b) Representation of a single base “flipping out of the stack,” showcasing a phenomenon known as DNA breathing. (c) Illustration of consecutive base-pairs breaking the H-bonds and opening simultaneously, referred to as DNA bubbles. (d-e) 3D surface plots highlighting the change in bubble intensity across varied lengths and base pairs (bps) for threshold value 1.5 under two conditions: P5 wild (d) and P5 mutant (e). (f-g) Average Coordinates profiles for AAV P5 wild (f) and mutant-promoter (g) sequences at individual base pairs, with the orange vertical block indicating nucleotide substitutions from AG to TC at the 50 and 51 positions (zero-indexed). (h) Average flipping profiles alongside the (i) corresponding q-factor for AAV P5 wild and mutant-sequence. For all the experiments, we set a minimum of 100 MCMC simulations using various initial conditions, setting the temperature to 310 Kelvin and employing 50,000 preheating steps followed by 80,000 post-preheating steps.

In living cells, the thermal fluctuations spontaneously induce local opening and closing of the DNA helix, referred to as DNA breathing, which can occur as a single base **flipping out of the stack** (base flipping) Figure 1b), or as a few **consecutive base-pairs breaking the H-bonds and opening simultaneously** (DNA bubbles) Figure 1c). Large amplitude bubbles describe local regions of DNA melting (denaturation).

Single base-pair opening/flipping, has been studied by tracking the exchange of protons from imino groups with water Guéron et al. (1987). DNA breathing is essential for DNA transcription, transcription factor (TF) binding, replication, enzymatic repair, and base pair methylation, Klimašauskas and Roberts (1995); Roberts and Cheng (1998); Choi et al. (2004); Dellarole et al. (2010); Fenwick et al. (2011); Alexandrov et al. (2012); Phelps et al. (2013),It was also observed that single base pair genetic variants, e.g., in the flanks of TF motifs) may influence DNA breathing dynamics and TF binding Alexandrov et al. (2011); Jablensky et al. (2012); Alexandrov et al. (2012); Duan et al. (2014).

The thermo-mechanical characteristics of DNA have been precisely described using a variety of models. Here We employ the mesoscopic Extended-Peyrard-Bishop-Dauxois (EPBD) model to examine DNA breathing dynamics. EPBD is an expansion of the Peyrard-Bishop-Dauxois nonlinear model of DNA and includes sequence-specific base-pair stacking potentials DNA breathing dynamics. The probabilities for local DNA openings obtained from the EPBD model are equilibrium properties of the underlying free energy landscape, and similar information can be obtained from various thermodynamical models, such as the Poland-Sheraga model.The parameters of the EPBD model have been derived from DNA melting experiments Ares et al. (2005); Alexandrov et al. (2009). The EPBD derived trajectories contain unique information about the relative **lifetimes of the DNA bubbles** which cannot be obtained by thermodynamical calculations. Another advantage of EPBD is its **single nucleotide resolution**.

By using EPBD simulations to locate DNA breathing hotspots, we were able to:(i) design (computationally) single-point mutations that silence breathing dynamics of transcription start sites (TSS), and to demonstrate (experimentally) that such mutations mutations also alter transcription, without affecting the TF binding Alexandrov et al. (2010); ii) design (computationally) mismatches, which enhance bubble formation, that can lead (experimentally) to bidirectional transcription initiation in the absence of basal TFs Alexandrov et al. (2009); (iii) design (computationally) base-pair substitutions or methylations, that change local DNA breathing and show (experimentally) that TF-DNA binding changes accordingly Nowak-Lovato et al. (2013). Further, EPBD simulations have indicated significant changes in the spatio-temporal characteristics of double-stranded DNA in the presence of a UV-dimer Blagoev et al. (2006) (which can be interpreted as an effective increase of the local temperature).

The EPBD modeling framework has been used to simulate the influence of non-ionizing terahertz (THz) radiation on DNA breathing and demonstrated Alexandrov et al. (2010) that, at sufficient exposure, DNA bubbles can appear through a nonlinear resonance mechanism. Such resonance By using EPBD simulations to locate DNA breathing hotspots, we were able to:(i) design (computationally) single-point mutations that silence breathing dynamics of transcription start sites (TSS), and to demonstrate (experimentally) that such mutations mutations also alter transcription, without affecting the TF binding Alexandrov et al. (2010); ii) design (computationally) mismatches, which enhance bubble formation, that can lead (experimentally) to bidirectional transcription initiation in the absence of basal TFs Alexandrov et al. (2009); (iii) design (computationally) base-pair substitutions or methylations, that change local DNA breathing and show (experimentally) that TF-DNA binding changes accordingly Nowak-Lovato et al. (2013). Further, EPBD simulations have indicated significant changes in the spatio-temporal characteristics of double-stranded DNA in the presence of a UV-dimer Blagoev et al. (2006) (which can be interpreted as an effective increase of the local temperature). The EPBD modeling framework has been used to simulate the influence of non-ionizing terahertz (THz) radiation on DNA breathing and demonstrated Alexandrov et al. (2010) that, at sufficient exposure, DNA bubbles can appear through a nonlinear resonance mechanism. Such resonance bubble formations may have a direct effect on transcription, replication, and DNA-protein binding, thus providing/suggesting a connection between THz radiation and biological function. Experimentally, it have been demonstrated that intense ultra-short THz pulses can indeed lead to non-thermal effects, including, modified gene expression profiles, gene-specific activation/repression, and changes in stem-cell differentiation Bock et al. (2010); Alexandrov et al. (2011, 2013); Titova et al. (2013); Perera et al. (2019); Shang et al. (2021). Importantly, based on the optical Kerr effect, it was shown that the interstrand H-bond modes (bubbles) of DNAare coherent delocalized/non-linear phonon modes in the THz range at physiological conditions González-Jiménez et al. (2016).

Experimental studies of looping of ultra-short DNA sequences revealed a discrepancy of up to six orders of magnitude between experimentally measured and theoretically predicted Jacobson-Stockmayer’s J-factors Vafabakhsh and Ha (2012); Alexandrov et al. (2016), suggesting that, in addition to the elastic moduli of DNA, the presence of local single-stranded “flexible hinges” in DNA (i.e., DNA bubbles) can assist the loop formation Yan and Marko (2004). Combining the Czapla-Swigon-Olson structural model of DNA with the EPBD model Alexandrov et al. (2017), (without changing any of the parameters of the two models), it was shown that the calculated J-factors of calculated J-factors of ultra-short DNA sequences are within an order of magnitude of the experimental measurements

## 2 Material and Methods

Our pyDNA-EPBD model utilizes Markov chain Monte Carlo (MCMC) protocol. The specific algorithm is summarized in Figure 1l). We use the standard Metropolis algorithm to produce an equilibrium state of the system: First, a base pair, *n*_0_, is selected at random, then a strand (left or right) is selected randomly and a new value of the variable 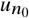 or 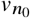 is proposed according to a thermal (at the given temperature) Gaussian distribution at this base pair. The proposed value is accepted according to the Metropolis probability, *P*: (a) *P* = 1 if the energy, *E*_*new*_, of the new configuration is lower than the energy, *E*_*old*_, of the old configuration; and, *P* = exp [(*E*_*old*_ – *E*_*new*_)] / *k* _*B*_*T*], if *E*_*new*_ *> E*_*old*_. The process is continued after thermal equilibrium is reached in the measurement phase. For each temperature, we performed several simulations (runs). In each of these runs, we compute the transfers opening profile *y*_*n*_. Subsequently, the displacements *y*_*n*_ of each base pair is recorded at every Δ*t* (intervals of selected MCMC steps). By conducting numerous runs, each with different initial conditions, we derived the average displacement/opening profile, denoted as ⟨*y*_*n*_⟩

One of the main characteristics of DNA breathing and DNA bubbles is the displacement, *y*_*n*_, (see Fig 1e) from equilibrium position for each base pair in the sequence. This displacement signifies the transverse stretching of the Hbonds between the complementary nucleotides. An advantage of the *y*_*n*_ profile is that it eliminates the need for window averaging typically required in thermodynamic calculations, thus making the average displacement/opening profiles sensitive to single base pair substitutions. The *y*_*n*_ profile can be efficiently calculated by MCMC simulations, yielding results equivalent to those obtained by averaging over Langevin dynamics trajectories Choi (2004). Below we introduce the main DNA breathing characteristics that our tool pyDNA-EPBD calculates:

### Average Coordinates

Refers to the averaged transverse displacements, ⟨*y*_*n*_⟩, i.e. the displacements, *y*_*n*_, averaged over the thermal fluctuations. The displacement profile, ⟨*y*_*n*_⟩, is one of the distinct properties of the DNA breathing characteristics, which quantifies the extent to which each of the base pairs of the DNA sequence is “open” in equilibrium, i.e., the extent to which the H-bonds between the bps are stretched due to the thermal fluctuations.

### Base Flipping Probability

Describes a particular kind of base movement in which at least one of the bases, in a base pair, flips out of the stack. The flipping exposes the base to the surrounding environment, which can be important for a variety of biological processes such as DNA repair, replication, and TF binding. The propensity of flipping characterizes this transition, by determining the fraction of disrupted Hbonds (openings) between complimentary nucleotides, i.e., the fraction of bps for which *y*_*n*_ exceeds a certain threshold distance, as a function of temperature.

### DNA Bubble Probability

Refers to extended DNA regions where the double helix unwinds and the two strands temporarily separate, due to the thermal motion. These transient opening or denaturation bubbles are part of the natural dynamics of DNA and can play a role in processes such as the initiation of transcription, replication and TF binding propensity. The probability for bubble existence, of a bubble begining at bp *n*-th, of length *l* bps and with a displacement *y*_*n*_ exceeding a given threshold (*thr*) (in Å) is a three dimensional (3D) tensor, *P*(*n, l, thr*), which can be computed as, Ref. Alexandrov et al. (2006),

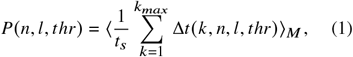

Here, ⟨·⟩_*M*_ denotes averaging over M runs, *t*_*s*_ is the total number of of MCMC simulation steps. The bubble duration Δ*t* (in MCMC steps) is a nonlinear function of its initial position (the beginning of the bubble), *n*, on its length, *l*, and amplitude threshold, *thr*.

### Dynamic Length

Since the average flipping profile represents the breathing propensity of each of the base pairs in a DNA region, we need a way to compare breathing profiles of two alleles that differ, e.g., due one or two SNPs. The dynamic length quantifies the statistical significance of the differences between two such flipping profiles. Specifically, the dynamic length, quantifies how many and precisely which base-pairs experience statistically significant changes caused by the introduced SNPs. We quantified the dynamic length by the concept of a “q-factor” as a function of the position of the base pair, which has been used to estimates the false discovery rate (FDR) in multiple stochastic simulations/experiments Lai (2017).

The average coordinates profile, (Figure 1(f)), the flipping probability profile, (Figures 1(g)), the 3D bubble probability tensor (Figures 1(d-e)), and the dynamic length profiles (Figure 1(h-i)), are key factors for understanding the DNA breathing. The corresponding details are provided in the supplementary material.

## 3 Usage

The pyDNA-EPBD tool is distributed under the 3-Clause BSD License, and it is compatible with Windows, MacOS, and Linux-based systems. pyDNA-EPBD is designed to work seamlessly in large-scale deployments, whether on high-performance computing clusters or cloud infrastructures like Amazon Web Services. Users can provide the input DNA sequence in common formats, such as FASTA or text file. Upon running simulations with specific parameters, pyDNA-EPBD generates the following characteristic outputs: the average coordinates profile, the flipping probability profile, the 3D probability bubble tensor, and the dynamic length profile. The details on hyperparameters, experimental configuration for reproducibility and analysis is provided in the supplementary material.

## Declarations

## Availability of data and materials

pyDNA-EPBD is Open Source Software published under the 3-Clause BSD License and can be found at https://github.com/lanl/pyDNA_EPBD.

## Authors’ contributions

AK and MB developed and implemented pyDNA-EPBD, and AK performed all validation experiments. BSA, KOR, ARB, and AU developed and validated the EPBD algorithm in ANSI C, which served as the base of pyDNA-EPBD implementation. MB, AS, and BSA consulted AK simulations. All authors wrote and edit the manuscript.

## Supplementary Materials A Implementation

To study DNA breathing dynamics, we used the mesocopic EPBD model, which is an extension of the original PeyrardBishop-Dauxois model Dauxois et al. (1993) that includes sequence-specific stacking potentials. A comment on the choice of model is perhaps appropriate, as many models have been used to study the mechanical properties of DNA. Most of them are purely thermodynamical models parameterized on the basis of measurements of equilibrium thermodynamical properties. The probabilities for local DNA opening obtained from the EPBD model are also equilibrium properties of the underlying free energy landscape, and essentially the same information can be obtained from various available thermodynamical models, such as the Poland-Sheraga model Richard and Guttmann (2004). However, it should be noted that the EPBD model is a dynamical model that is strongly nonlinear and admits breathing solutions, which constitute transient but relatively long-lived openings of the double helix, that are interconnected with the local bending propensity. The EPBD derived trajectories contain the information about the lifetimes of the DNA transient openings (bubbles). This type of information cannot be obtained by purely thermodynamical calculations. Explicit accounting of the dependence on the solvent conditions such as salt, temperature, as well on the twist of the DNA, can lead to long-lived bubbles with enhanced lifetimes. Another advantage of the EPBD model is its single nucleotide resolution. With a thermodynamical model, the calculation of a property profile typically requires window averaging over 100–500 bps, which limits one’s ability to distinguish the property profiles of two closely related sequences. In contrast, window averaging is not used in EPBD calculations, and as a result, the effects of even single bp changes can be readily determined.

The EPBD model is a quasi-two-dimensional nonlinear model that describes the transverse opening motion of the complementary strands of double-stranded DNA, while distinguishing the two sides (right-*u*_*n*_ and left-*v*_*n*_) of the DNA double strand. The Hamiltonian potential surface *V*_*EPBD*_ of the EPBD model is defined by the summation of the Morse Potential (first part) and stacking energy term (second term) of the two neighboring bps at every bp of the input DNA fragment, see Eqn. 2.

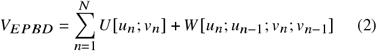

The Morse potential incorporates the Hydrogen bonds connecting two bases belonging to opposite strands, the repulsive interactions of the phosphates, and the surrounding solvent effects. The parameters D and a in this term denote the nature of the *n*-th bp, i.e. A–T or G–C, see Eqn. 3.

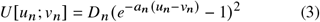

The stacking energy potential is defined between two neighboring bps using their status variables *y*_*n*_ and *y*_*n* −1_ as in Eqn. 4 which is defined by the distance of the bp, *y*_*n*_ = *u*_*n*_ −*vn*. The *K* term determines the coupling constants between the left and right two neighboring bases.

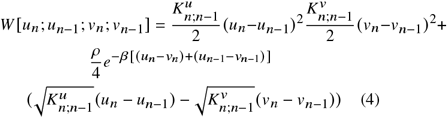

MCMC methods, for sampling probability distribution, constructs Markov chain with the target distribution. By recording states of the chain, one can obtain samples of a desired distribution. The more the number of samples, the more the samples distribution gets closer to the actual distribution. The MCMC protocol used in pyDNA-EPBD model is similar to what Ares *et. al* Ares et al. (2005) applied to understand the bubbles activity in DNA melting.

To calculate the average displacement or opening profile for a given DNA sequence at a specific temperature, we utilized the pyDNA-EPBD model in our MCMC simulations as detailed in previous literature, summarized in Algorithm 1 and in Figure 2. The standard Metropolis algorithm was applied to generate an equilibrium state in each simulation run. Subsequently, the displacements *y*_*n*_of each base at selected time intervals were recorded. By conducting numerous simulation runs, each with different initial conditions, we derived the average displacement/opening profile, denoted as *y*_*n*_, for the DNA sequence under investigation.

**Fig. 2:**
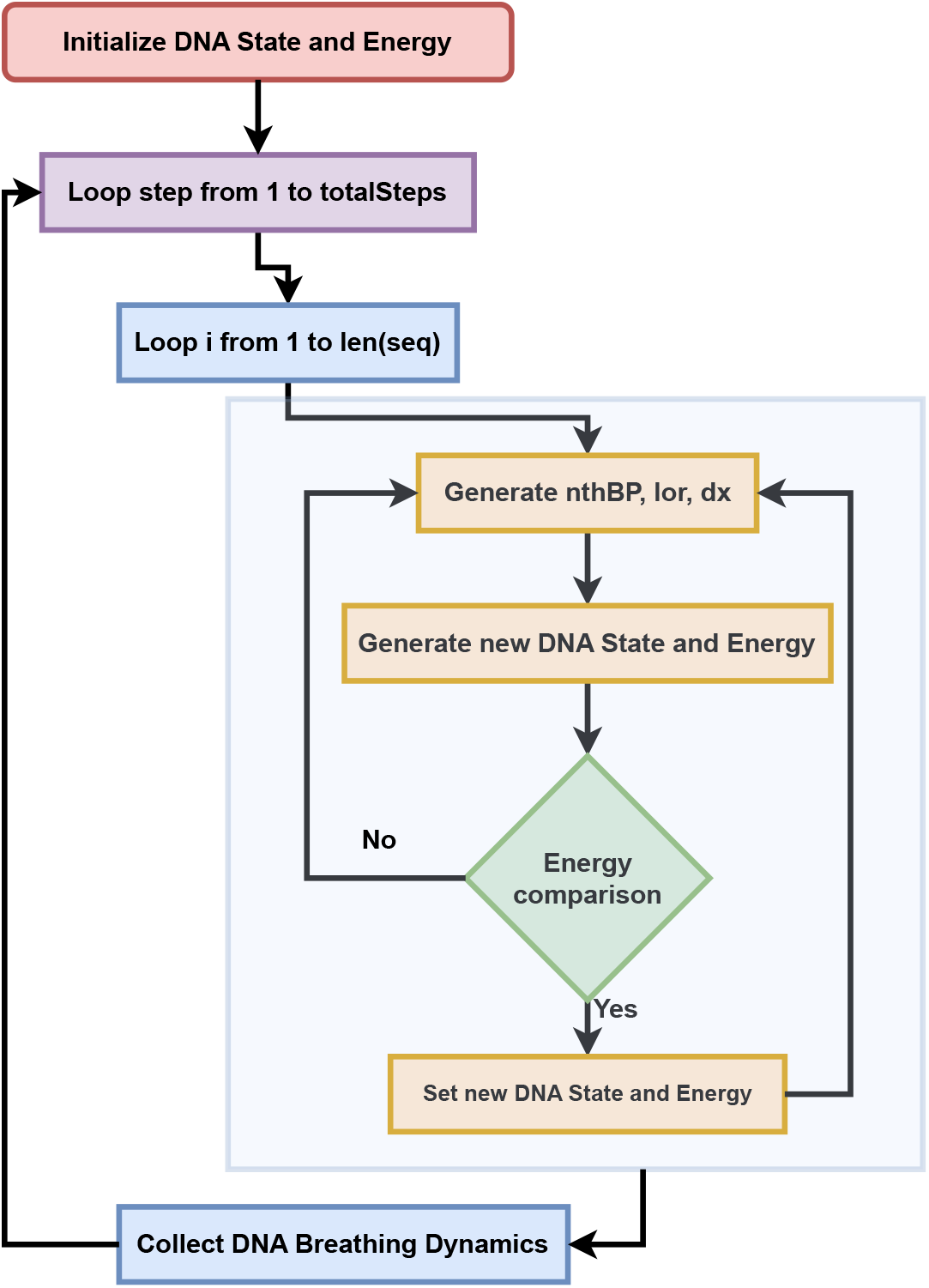
Schematic representation of the Metropolis-Hasting MCMC Algorithm for DNA State Transitions in pyDNA-EPBD

A single step in the simulation involves selecting a random base-pair to do random displacement at the left or right base. Based on the displacement occurred, new DNA state and corresponding energy is calculated using Morse-potential (Eqn. 3) and stacking energy (Eqn. 4) in the EPBD model (Eqn. 2). The new DNA state is updated if the energy tends to an equilibrium state or the energy change due to this random displacement satisfies a Metropolis criteria. The criteria is defined such that the energy change due to the random displacement follows a simulation temperature scaled log-uniform distribution.

### Algorithm 1: Metropolis-Hasting MCMC algorithm

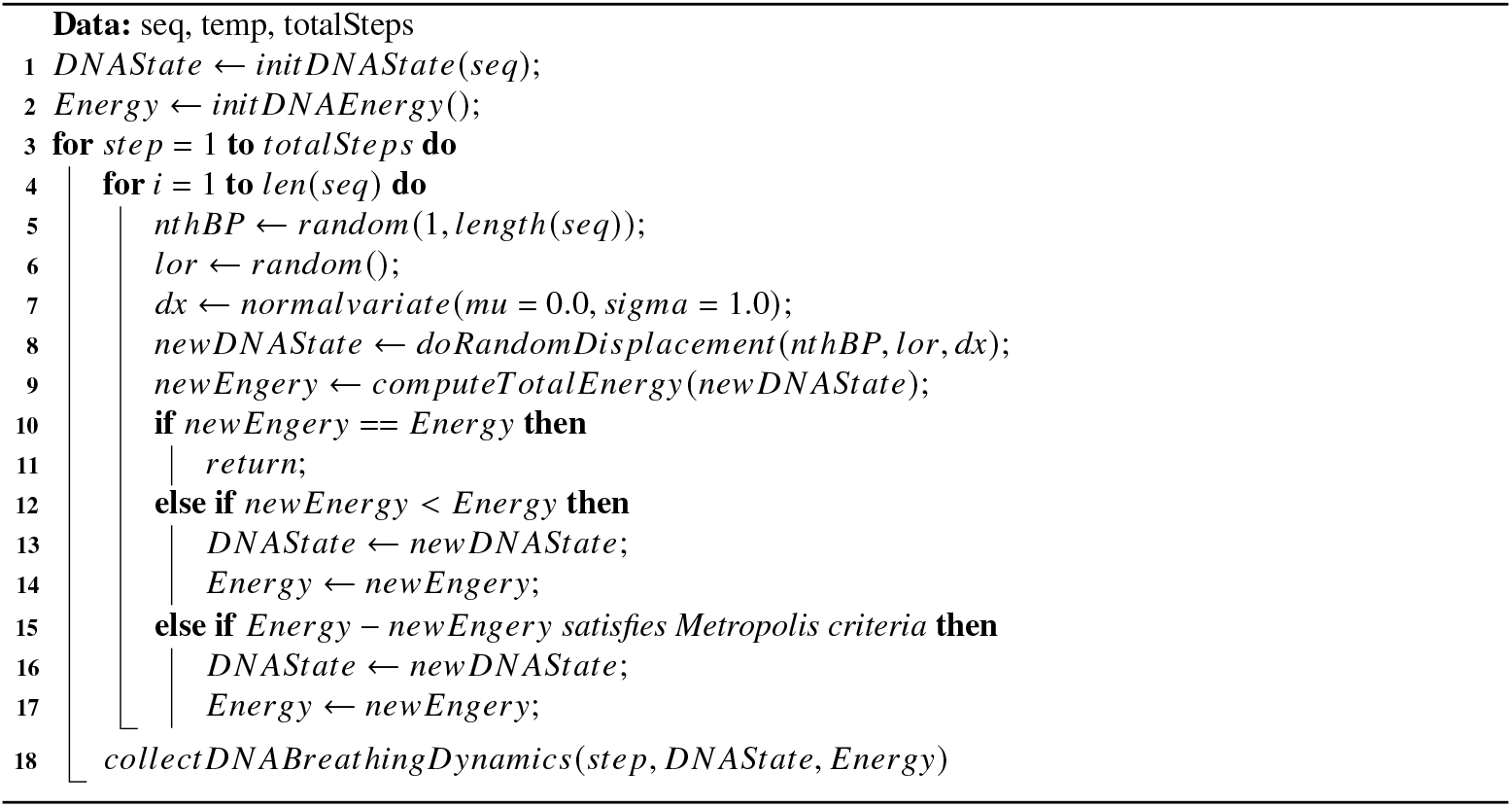

Figure 3 shows an example distribution of the chosen Metropolis criteria at 310.0 Kelvin. 130, 000 samples are randomly generated from the criteria for demonstration. The violin plot shows the box-and-whisker plot (red-ish box with vertical caps) along with the kernel density estimate (KDE) (blue area). The KDE shows the continuous probability density using a Gaussian kernel which is skewed towards zero, indicating that a new DNA state will be selected during the MCMC simulations if the energy change is small. This criteria ensures that the simulation does not get stuck in the local energy minima, instead, it will explore the energy landscape to reach a possible global minima. In particular, the box and whisker plot shows that 75% of the samples are within range from 0 to 51.78 Kelvin (up to third quartile) which gives the empirical evidence of such exploration. In essence, this Metropolis criteria ensures that the simulation reaches an equilibrium state, possibly a global minima, by examining enough Markov samples in the energy landscape.

**Fig. 3:**
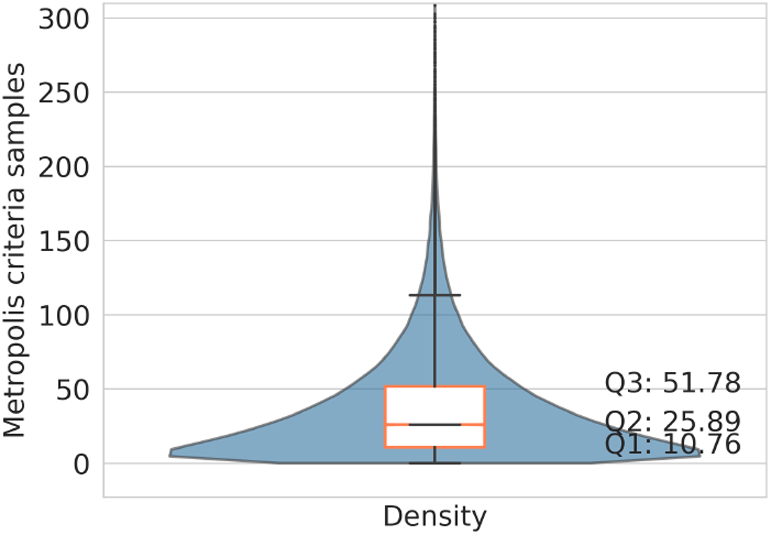
Continuous probability (Kernel density estimate) distribution of the Metropolis criteria (violin plot, blue area). Y-axis shows the spread of the generated 130, 000 samples (simulation temperature scaled in Kelvin) and x-axis gives the density. The box-and-whisker plot facilitates the distribution in quartiles. Both demonstrates pyDNA-EBPD’s capability of finding an equilibrium DNA state by exploring its energy landscape.

## B Results and Discussions

First, we provide the specifics of datasets and the simulation configurations used throughout the study. Subsequently, we discuss the DNA breathing dynamics, such as the coordinates of bps, the flipping of bases, the formation of bubbles, and the q-factor. We also detail an utility of the breathing characteristics for Transcription Factor (TF)-DNA binding specificity prediction. Finally, we analyze runtimes of our pyDNA-EPBD model as a function of the number of bps consisting each DNA sequence.

### B.1 Datasets

This section briefly discusses the datasets utilized throughout the study, necessary to understand different utilities and perspectives of our pyDNA-EPBD model.

#### Adeno-associated virus (AAV) P5 promoter

Experimental data suggest that a spontaneous double strand DNA (dsDNA) separation at the transcriptional start site is likely to be a requirement for transcription initiation in several promoters Alexandrov et al. (2009). To study this phenomena using our pyDNA-EBPD model, we utilize the strand separation (bubble) dynamics of 77-bp-long AAV P5 promoter (wild-type (wt) sequence: “GCGCGTGGCCATTTAGGGTATATATG-GCCGAGTGAGCGAGCAGGATCTC-CATTTTGACCGCGAAATTTGAACGGCGC”). We also study the control non-promoter sequence (77 bp) (mutant-type (mt) sequence: “GCGCGTGGCCATTTAGGGTATATATGGCC-GAGTGAGCGAGCAGGATCTCCGCTTTGAC-CGCGAAATTTGAACGGCGC”), part of the published sequence for the human collagen intron (NW_927317), to do comparative analysis of the impact of the mutation .

#### DNA fragments

The propensity of cyclization determines the intrinsic bendability of a DNA fragment which may influence a myriad number of essential cellular mechanisms. Previously Alexandrov *et. al* Alexandrov et al. (2016) studied the impact of cyclization rates based on DNA structural parameters of 86 DNA fragments of length range from 57 to 325 bp (excluding flank sequence “GC” on both sides in our study) with experimentally determined cyclization factors.

#### Seven SNPs

To study the functional impact of single nucleotide polymorphisms (SNPs), we studied seven SNPs with the original sequence and allele modified sequence at and around 100 bp of the mutation site. The studied reference SNP ids (rs-ids) are reported in Section B.6.

### B.2 pyDNA-EPBD simulation

Given an input DNA sequence, following Algorithm 1, we run at least 100 MCMC simulations with different initial conditions. The temperature is set to 310.0 Kelvin with 50, 000 and 80, 000 preheating and post preheating steps, respectively, throughout the studies. Each random displacement is performed by selecting a random base-pair’s updated coordinate cutoff threshold of 25 Å. The breathing dynamics are recorded throughout the post preheating steps at different runs as the samples of the generated Markov chain. The distribution of the samples is computed over the independent runs and normalized over the number of post preheating steps as the final breathing characteristics. The next sections discusses the properties of the simulation outputs as the DNA breathing dynamics in light of capturing several DNA breathing profiles.

When utilizing the pyDNA-EPBD model for analysis of short DNA sequences, the addition of flanking sequences can play a critical role. This is primarily because the biophysical properties of DNA often extend beyond the confines of the sequence of interest and are influenced by the broader bp context. The biophysical traits of a short sequence, in isolation, may not fully encapsulate these sequence-dependent effects. Thus, by including flanking sequences, we are better able to simulate the native context of the DNA sequence, enhancing the accuracy of the model’s outputs. Another key factor is the potential distortion of the overall characteristics of short sequences by terminal bps. Given their significant proportion in relation to the total length of the sequence, the terminal bps can have a disproportionate influence on the sequence’s characteristics. Therefore, by integrating flanking sequences, we can moderate these boundary effects, ensuring a more representative portrayal of the internal bps within our sequence of interest. Moreover, the pyDNA-EPBD model imposes algorithmic requirements that specify a minimum sequence length. Failure to meet this threshold can lead to inaccurate results. Here, flanking sequences serve a crucial role by augmenting the sequence length to satisfy the model’s prerequisites, thereby ensuring the production of reliable features. However, the necessity for flanking sequences diminishes as the length of the sequence increases. Long sequences are typically sufficient to fulfill the model’s computational requirements, and the influence of the terminal bps on the overall sequence characteristics becomes negligible, thereby reducing the need for additional flanking sequences.

### B.3 Breathing dynamics: Average coordinate distance profile

Alexandrov *et. al* Alexandrov et al. (2009) revealed that the simulation distribution matches the dramatic difference in the transcriptional activity of the promoters. An example showcase is demonstrated for AAV P5 wildand mutantpromoter sequences. Following the same analysis, we compute the average coordinate distance profile (*y*_*n*_) for the AAV P5 promoters using our pyDNA-EPBD model. Figure 4 (left panel) shows the average displacements at-and-around bp 50 of the AAV P5 promoter (blue) and a transcriptionally silent AG to TC mutant (red). This result is visually identical as previously reported in Alexandrov et al. (2009). One can utilize the average displacement magnitude in the double helix width as a functional effect on the transcription-factor binding affinity. Moreover, we report the average coordinate profile for 86 different DNA sequences ranging from 61 to 329 bp (including the flanks) in link1.

**Fig. 4:**
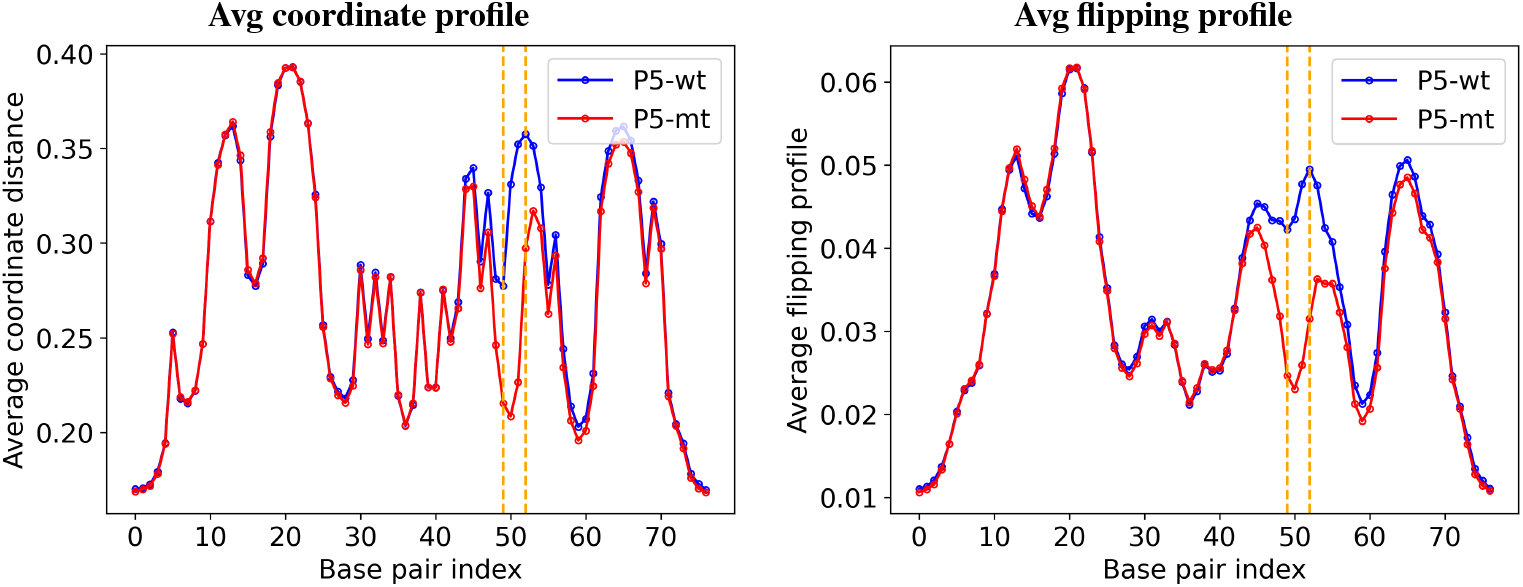
Average coordinate distance (left) and flipping (right) profile for AAV P5 wildand mutant-promoter sequences at each BP. The vertical block (orange) shows the mutation of a nucleotide bps substitutions from AG to TC at 50 and 51 positions (zero-indexed).

### B.4 Breathing dynamics: Average flipping profile

DNA breathing can expose nucleotide base openings for interaction with proteins, which can be favorable depending on the mechanism of TF-DNA binding Nowak-Lovato et al. (2013). These openings effectively correspond to the base flipping at various sites at physiological temperature and pH, where base flipping gives the base-pair mismatches.

We computed the average flipping profile using our pyDNA-EPBD model as the probability of a base-pair being flipped throughout the simulation steps. A basepair is considered flipped if the coordinate distance of that base-pair is no less than a threshold (in Å). We collected flipping profiles at five different thresholds from 0.707106781186548 Å to 3.53553390593274 Å, inclusive, with step size 0.707106781186548. Note that the floating point precision is crucial for obtaining accurate distribution. Figure 4 (right panel) demonstrates example flipping profiles for AAV P5 wild- and mutant-promoter sequences at threshold 1.414213562373096 Å. It shows that the transcriptionally silent AAV P5 mutant is less prone to bp openings at-and-around mutation position at certain threshold compared with its counter wild P5 promoter. Thus, average flipping profile is subjected to study more profoundly to understand the functional activity of the SNPs. We also provide the average flipping profile for 86 DNA nucleotide sequences at 1.414213562373096 Å threshold, which can be found in link2.

### B.5 Breathing dynamics: Bubbles

We collected bubbles using 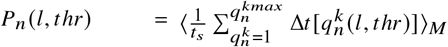 from minimum of three bps to maximum of 20 bps incrementally by applying our pyDNA-EPBD simulation tool at 20 different thresholds from 0.5 to 10.0 Å with step size 0.5. Figure 5 demonstrates the bubbles for AAV P5 wild- and mutant-promoter sequences on the left and right panel, respectively. Each row compares the bubbles with its counterparts at a certain threshold.

**Fig. 5:**
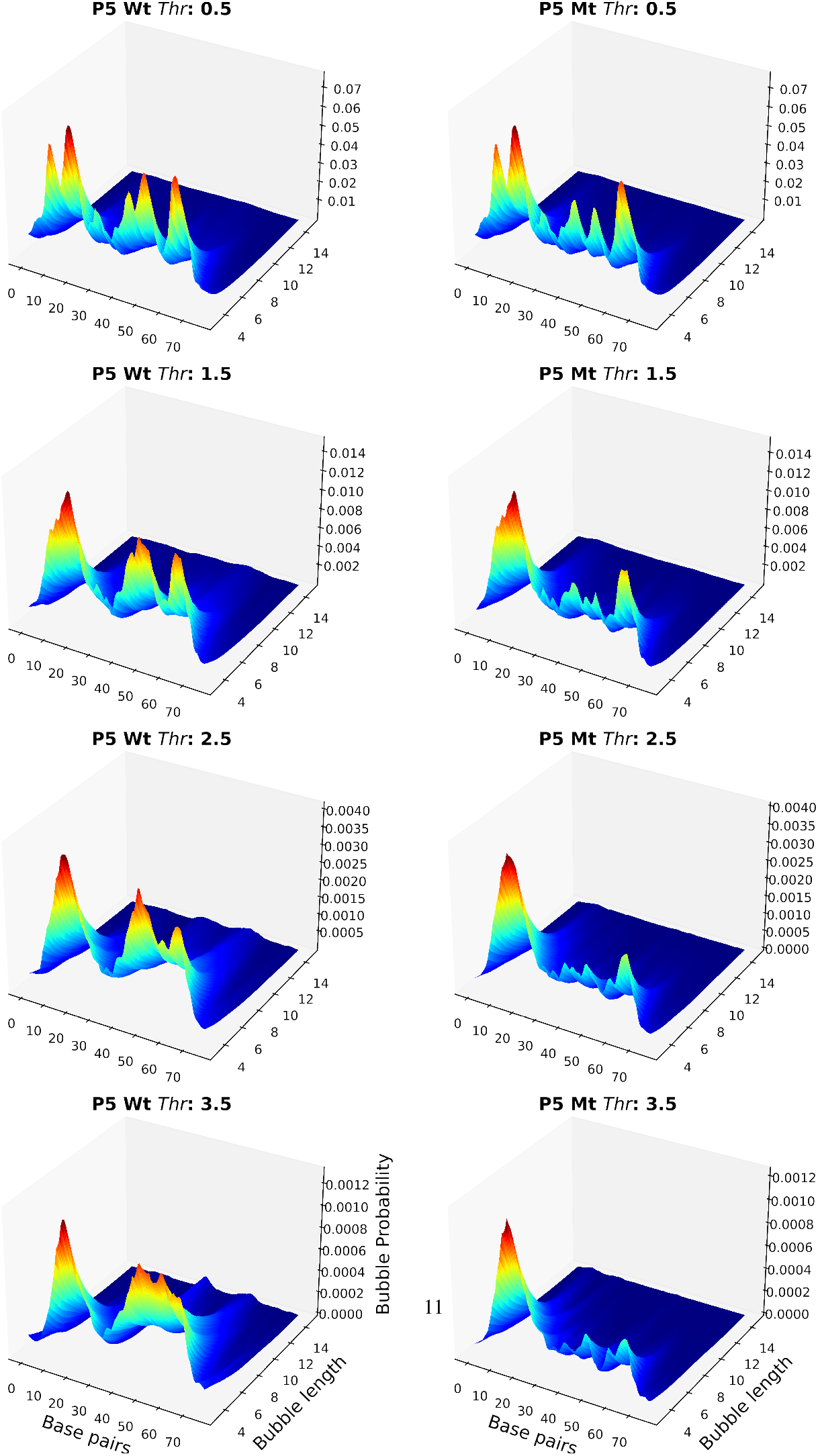
3D surface plots representing the change in bubble intensity across various bubble lengths and bps. Each row represents different threshold values for two different conditions: P5 wild (left panel) and P5 mutant (right panel).

In the first row, we can observe overall peak distribution when a small bubble detection coordinate distance threshold (*thr* = 2.5 Å) is considered. As the threshold got increased, bubbles are created at very different locations in AAV P5 wild- and mutant-promoters. This confirms previous findings in Choi (2004); Alexandrov et al. (2006, 2009) that a dramatic difference in the bubble probability at the mutation site in those two sequences has been occurred, matching the dramatic difference in the transcriptional activity of the promoters. Another important observation is that bubbles are only stretched over small number of nucleotide bps, which provides evidence of spontaneity of opening and re-closing of the double stranded DNA. Furthermore, this phenomena has previously been observed by Peyrard Peyrard (2004) in the early work of analyzing dynamical properties in DNA where only the bubbles with small stretching length are found when the simulation temperature is set to less than the denaturation temperature. Since we study all the simulations in this work at 310.0 Kelvin, we only observe small breathing window. One can also observe bubble denaturation using thermal bath by increasing simulation temperature.

### B.6 Breathing dynamics: Q-factor

Sequence-dependent DNA breathing dynamics from the nonlinear EPBD model Alexandrov et al. (2009) has been successfully used to demonstrate the effect of SNPs on TF binding via DNA breathing. While the breathing profile or flipping profile represents the average probability of DNA openings at each bp in the region, the MCMC simulation induces statistical errors in the distribution of the breathing characteristics. In Figure 6 (left panel), we show the average flipping profile with 95% confidence interval (shaded region). Thus, here we propose a novel dynamical feature named q-factor to compare the statistical differences in the breathing distribution at each bp between a SNP (wild and mutant pair). First we collect the flipping profile using our pyDNA-EPBD model for a SNP. At each bp we quantify a distance between two Markov distributions of the original sequence and allele modified sequence. We compute a statistical test known as Kolmogorov Smirinov2 test and scale the distances into negative logarithm (base 10). In Figure 6 (right panel), we report the q-factor for AAV P5 wild- and mutant-promoter. Clearly, the q-factor at the mutation gives a functional effect measurement which was not previously readily available.

**Fig. 6:**
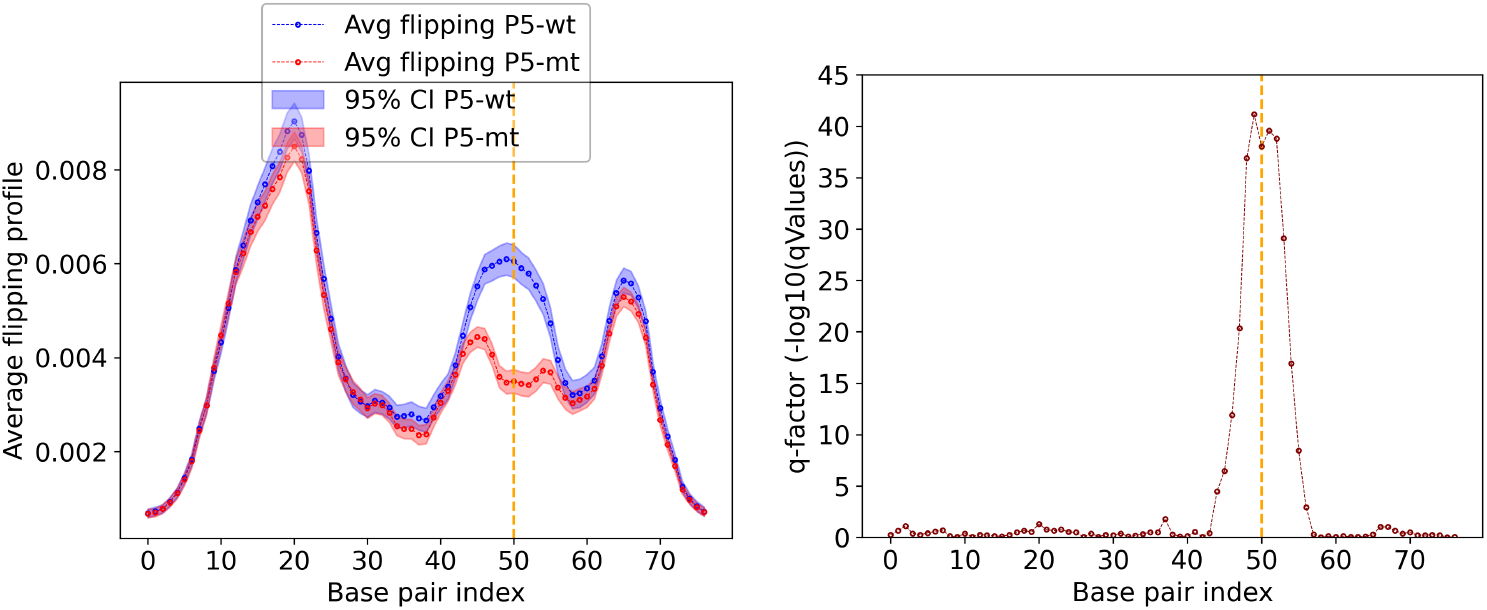
Average flipping profile with corresponding q-factor for AAV P5 wild- and mutant-sequence.

To summarize the effect of a SNP on DNA breathing in the 100 base-pair region, we computed the q-factor scores of three tested SNPs of PIQ-predicted TF footprints. These SNPs likely affect TF binding and have higher scores when compared to the four control SNPs that are not adjacent to PIQ-predicted TF footprints. As an example, in Figure 7 rs12946510, known to affect TF binding, significantly affects the DNA breathing of 11 base-pair around the SNP. In contrast, rs72974222, which is not adjacent to a PIQ-predicted TF footprint, weakly affects the DNA breathing of 5 base-pair around the SNP. In Figure 7 (bottom panel), we summarize the q-factor scores at the mutation site for seven SNPs and P5 promoter. As the figure clearly suggests, the boundary between PIQ-predicted TF footprint and controls SNPs is easily distinguishable. However, it requires an extensive study to draw a cutoff line which we deem as one of the future works.

**Fig. 7:**
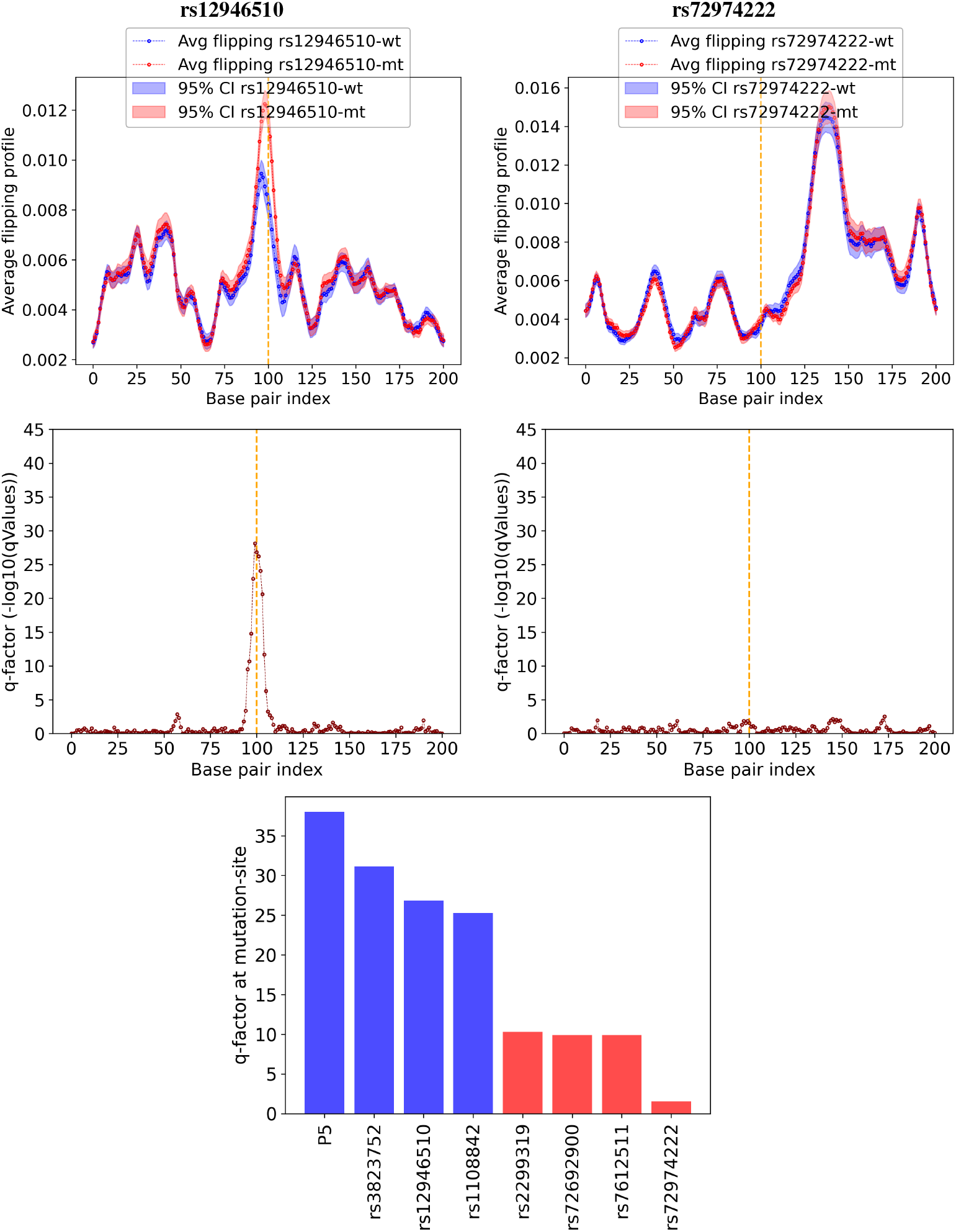
Average flipping profile with the corresponding q-factor for P5 two SNPs (rs12946510 and rs72974222 on the left and right panel). On the bottom panel, q-factor values for P5 and three PIQ-predicted TF footprint and four controls SNPs.

## Footnotes

1 Availability and implementation: pyDNA-EPBD is supported across most operating systems and is freely available at https://github.com/lanl/pyDNA_EPBD. Extensive documentation can be found at https://lanl.github.io/pyDNA_EPBD/. Contact

